# Individual heterogeneity and capture-recapture models: what, why and how?

**DOI:** 10.1101/120683

**Authors:** Olivier Gimenez, Emmanuelle Cam, Jean-Michel Gaillard

**Affiliations:** CEFE UMR 5175, CNRS, Université de Montpellier, Université Paul-Valéry Montpellier, EPHE, 1919 Route de Mende, 34293 Montpellier Cedex 5, France.; Evolution et Diversité Biologique, UMR 5174, CNRS-ENSFEA-IRD, Université Toulouse 3 Paul Sabatier – Bâtiment 4R1, 118 route de Narbonne, 31062 Toulouse Cedex 9, France.; Laboratoire de Biométrie et Biologie Evolutive, UMR 5558, Université de Lyon, Université Lyon 1, 69622 Villeurbanne, France.

**Keywords:** actuarial senescence, Arnason-Schwarz model, Cormack-Jolly-Seber model, frailty, hidden Markov models, individual covariates, life-history trade-offs, mark-recapture models, mixed models, mixture models, multievent models, multistate models, random-effect models, survival estimation

## Abstract

Variation between and within individuals in life history traits is ubiquitous in natural populations. When affecting fitness-related traits such as survival or reproduction, individual heterogeneity plays a key role in population dynamics and life history evolution. However, it is only recently that properly accounting for individual heterogeneity when studying population dynamics of free-ranging populations has been made possible through the development of appropriate statistical models. We aim here to review case studies of individual heterogeneity in the context of capture-recapture models for the estimation of population size and demographic parameters with imperfect detection. First, we define what individual heterogeneity means and clarify the terminology used in the literature. Second, we review the literature and illustrate why individual heterogeneity is used in capture-recapture studies by focusing on the detection of life-history trade-offs, including senescence. Third, we explain how to model individual heterogeneity in capture-recapture models and provide the code to fit these models (https://github.com/oliviergimenez/indhet_in_CRmodels). The distinction is made between situations in which heterogeneity is actually measured and situations in which part of the heterogeneity remains unobserved. Regarding the latter, we outline recent developments of random-effect models and finite-mixture models. Finally, we discuss several avenues for future research.

## Introduction

Individual variation is at the core of the evolution of traits by the means of natural selection and exists within any population of living organisms. Individual variation occurs in virtually all traits, including fitness components such as reproduction and survival (Clutton-Brock 1988, Newton 1989). However, the amount of individual variation in a given trait in a given population varies a lot both within and across species. Between-individual differences in phenotypic attributes such as age (Caughley 1966, Emlen 1970), sex (Short and Balaban 1994), body mass (Sauer and Slade 1987), or personality (Dingemanse and Dochtermann 2013), in genotype (Coulson et al. 2011), in habitat use or habitat selection such as home range size or quality (Mcloughlin et al. 2007), or in prey selection (Estes et al. 2003) have all been reported to affect most life history traits. More recently, both current and early-life environmental conditions encountered by individuals throughout their lives have been shown to generate individual differences in life history traits (Douhard et al. 2014, Berger et al. 2015).

The potential role of individual heterogeneity in terms of population ecology has been pointed out more than 30 years ago (Lomnicki 1978, Johnson et al. 1986) and repeatedly reported since (Bolnick et al. 2011, Kendall et al. 2011). Thanks to the increasing availability of high quality data collected during long-term individual monitoring of vertebrate populations (Clutton-Brock and Sheldon 2010), assessing the magnitude of individual heterogeneity, identifying its origin, and quantifying its consequences has become a specific objective in many population studies.

From these studies (reviewed in Table 1), we can envisage three broad patterns of individual heterogeneity when considering a set of life-history traits, e.g. demographic parameters, independently of any methodological approach used to model these demographic tactics.

**Table 1.**
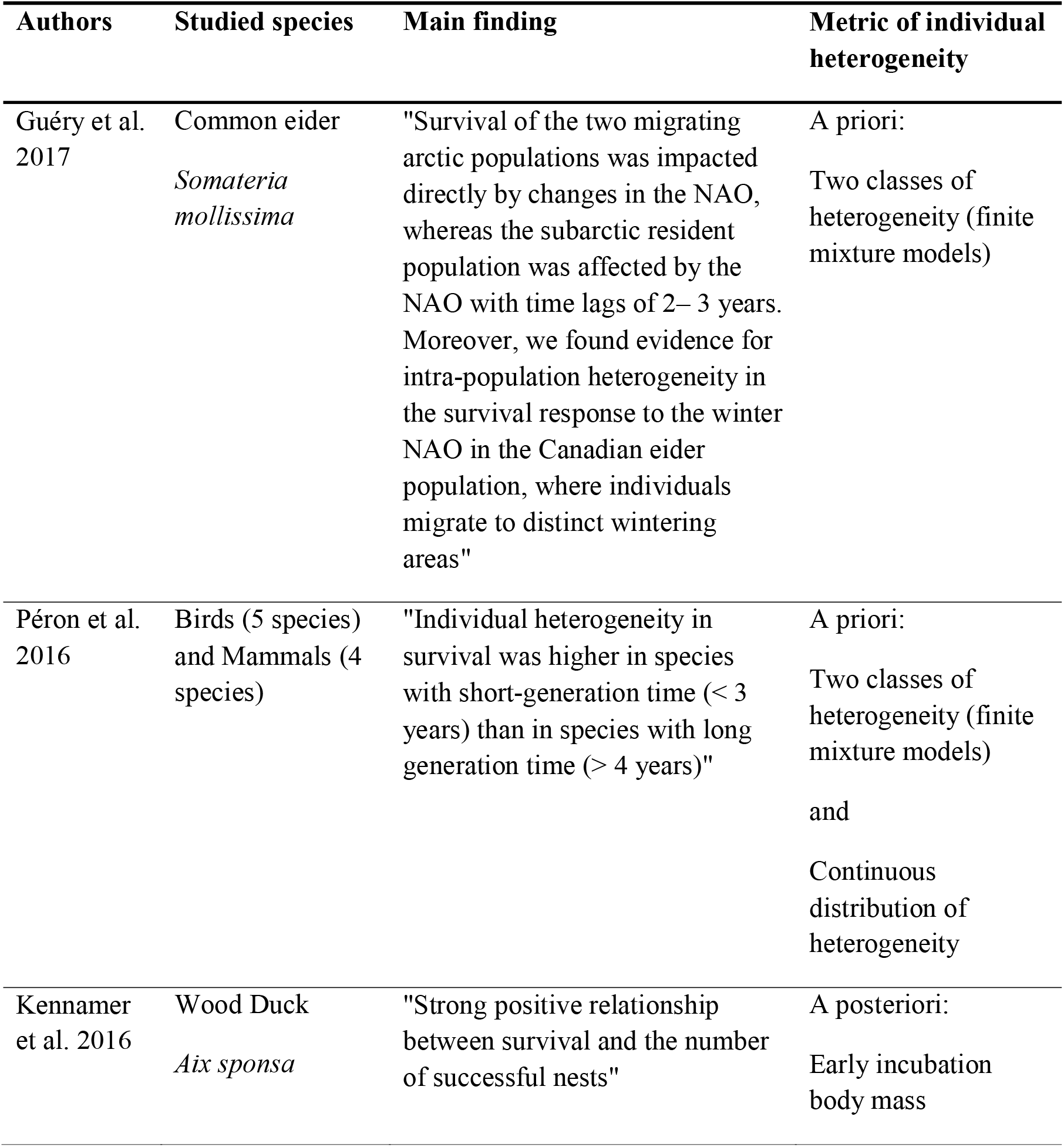
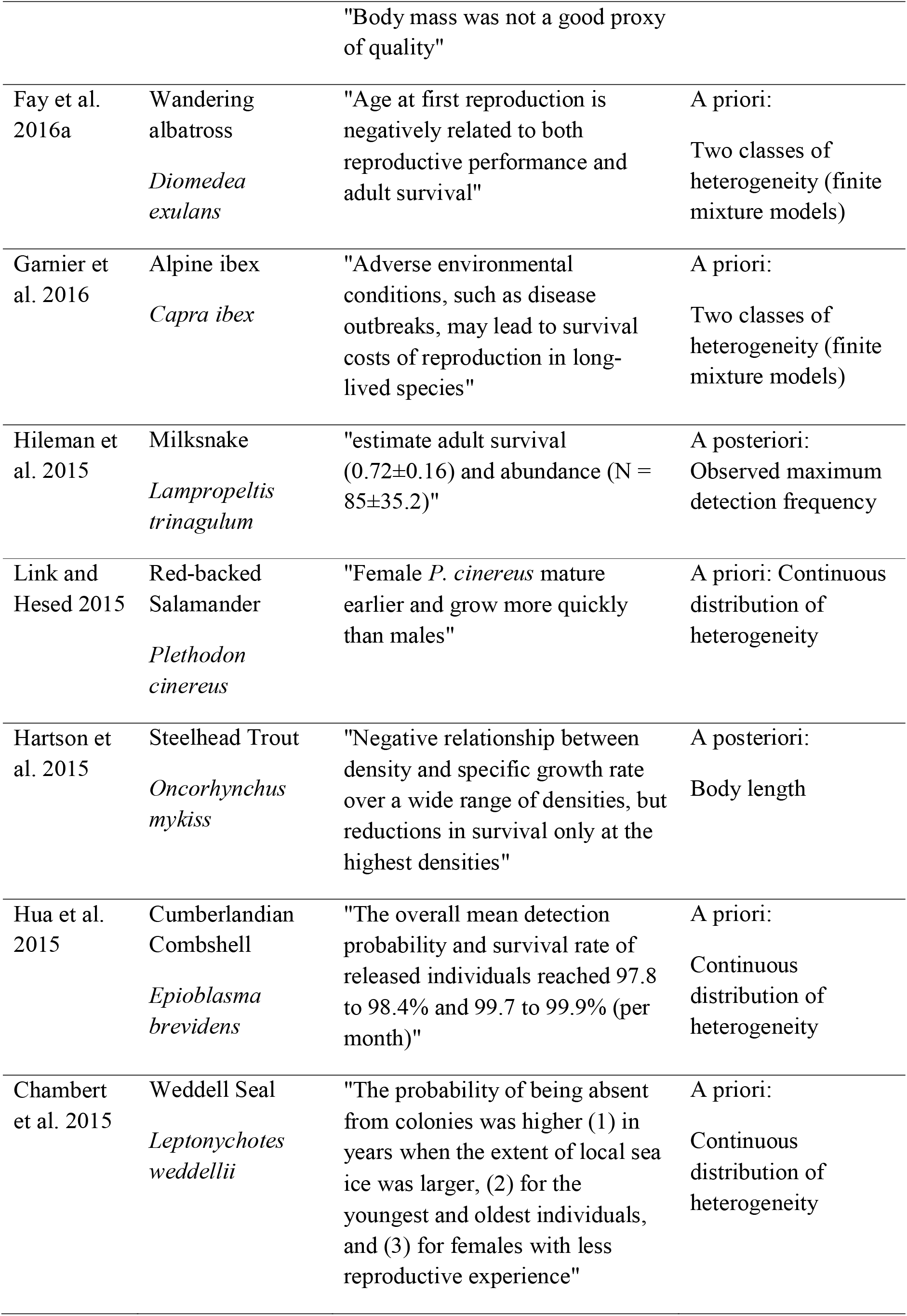
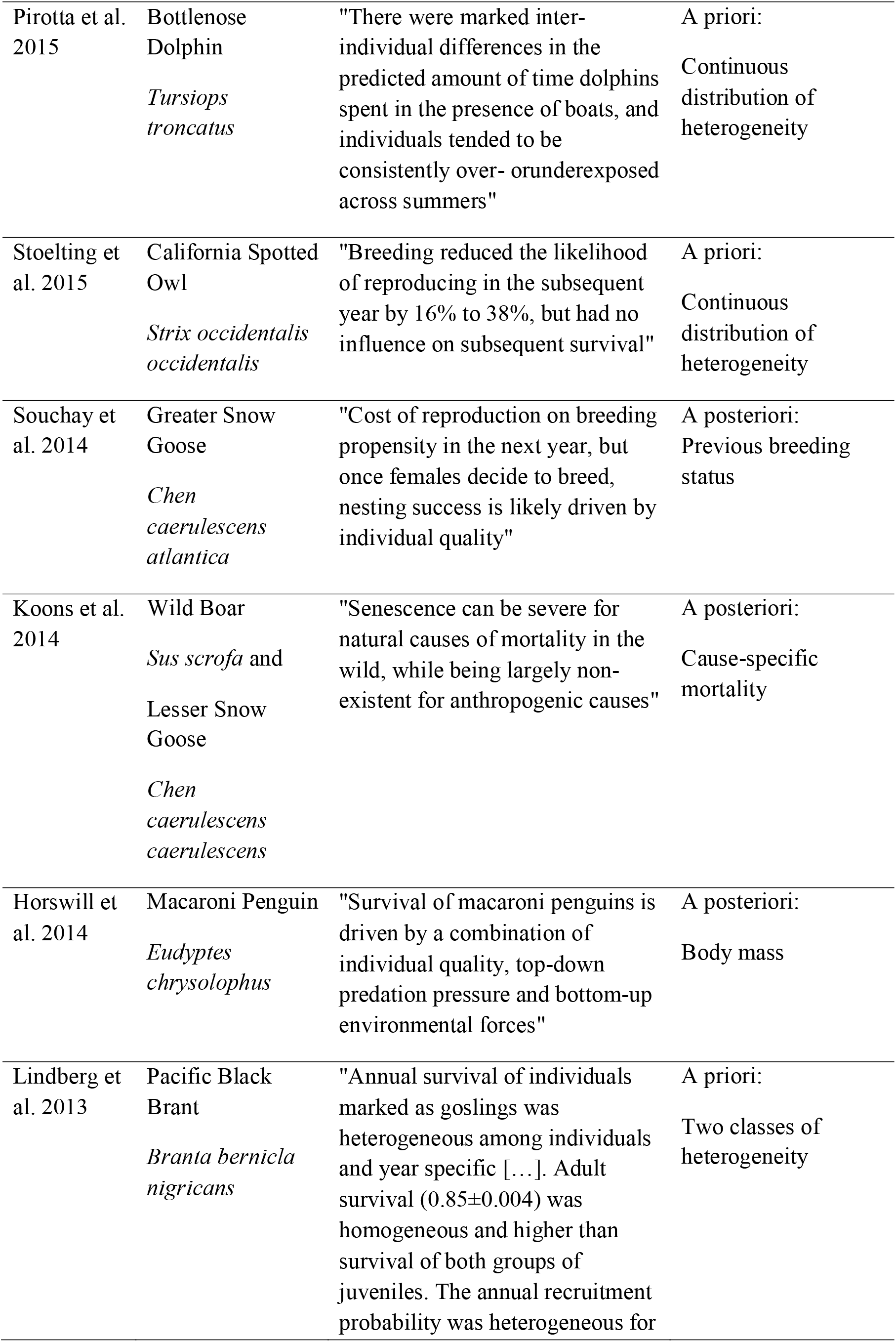
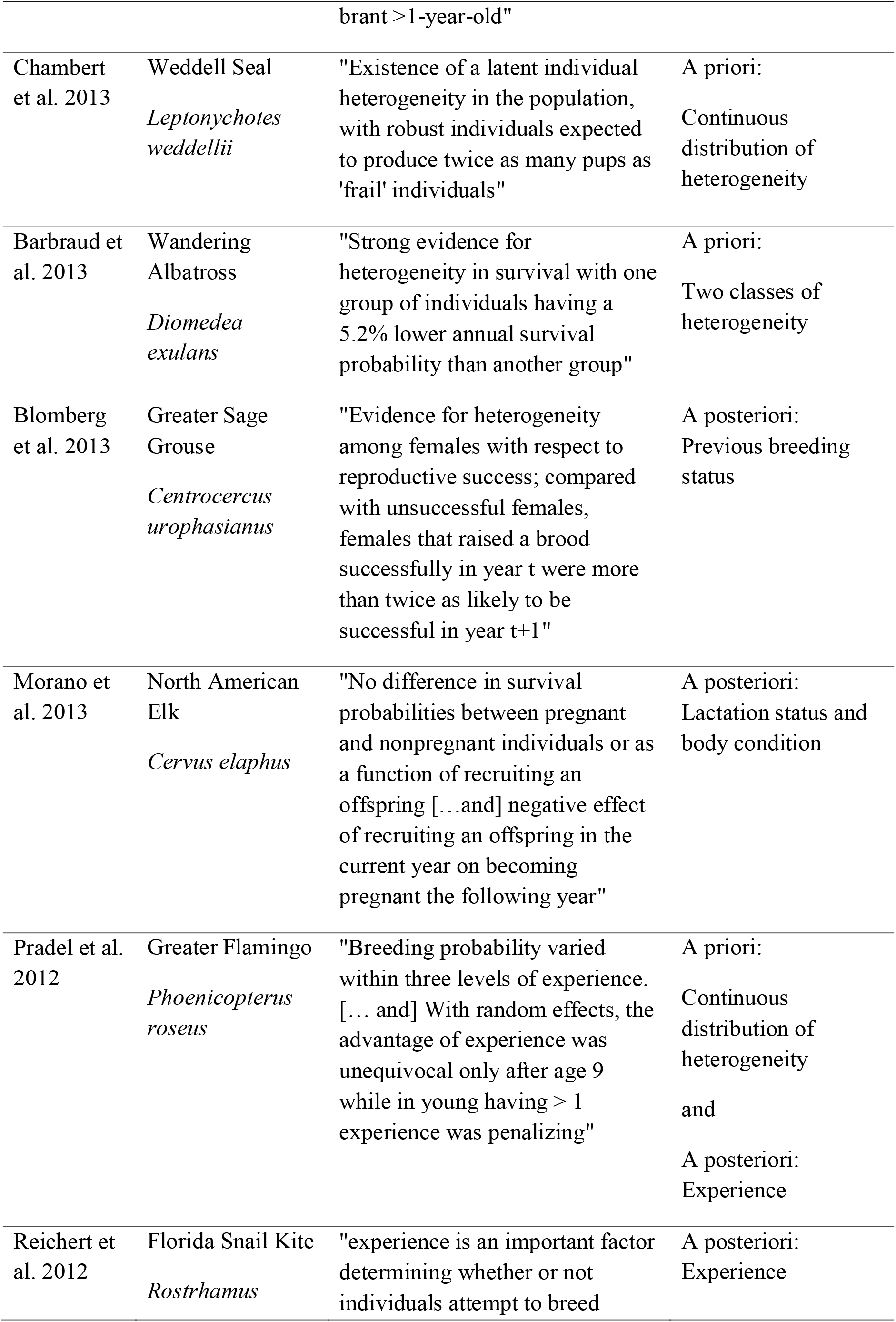
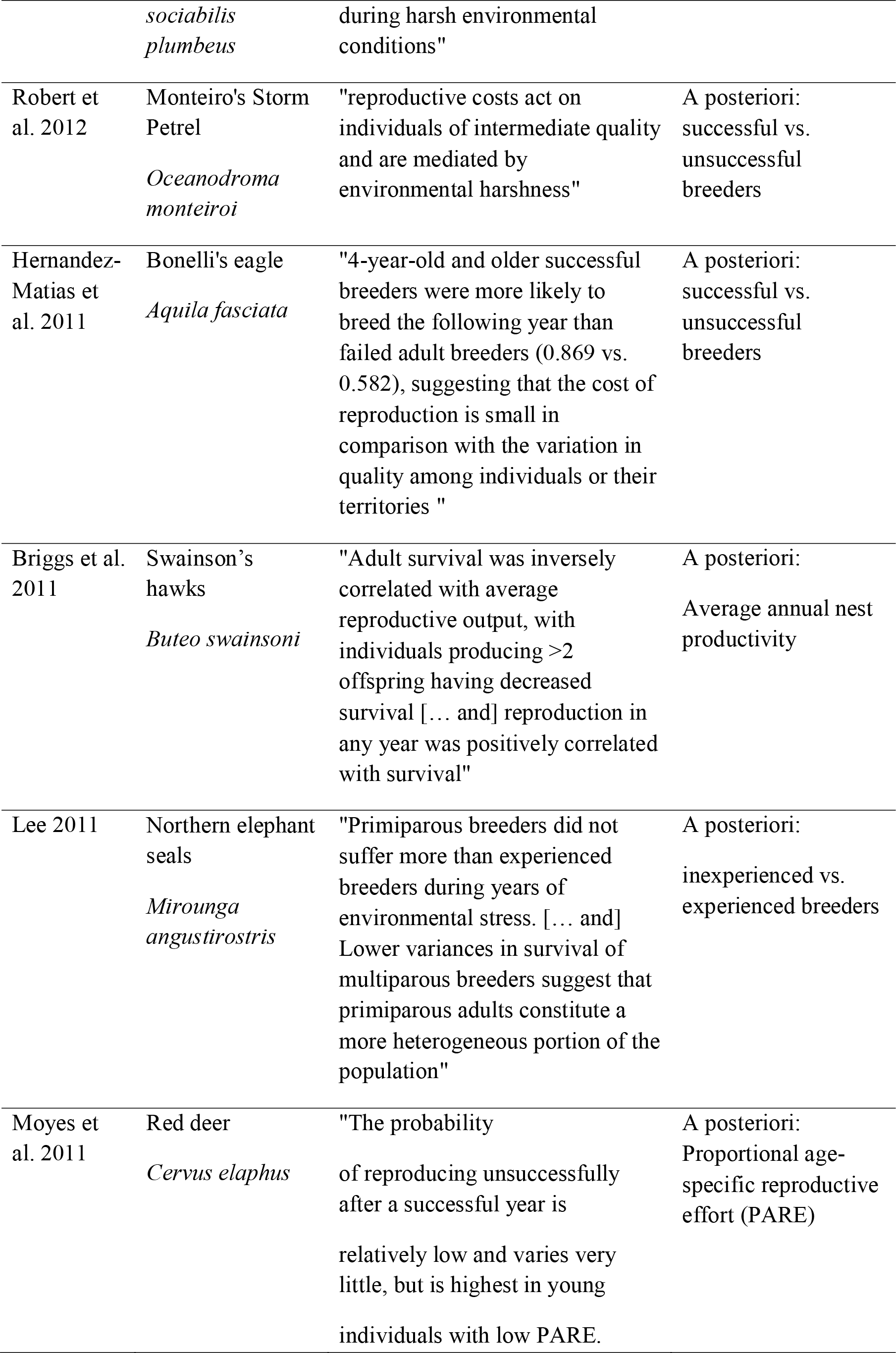
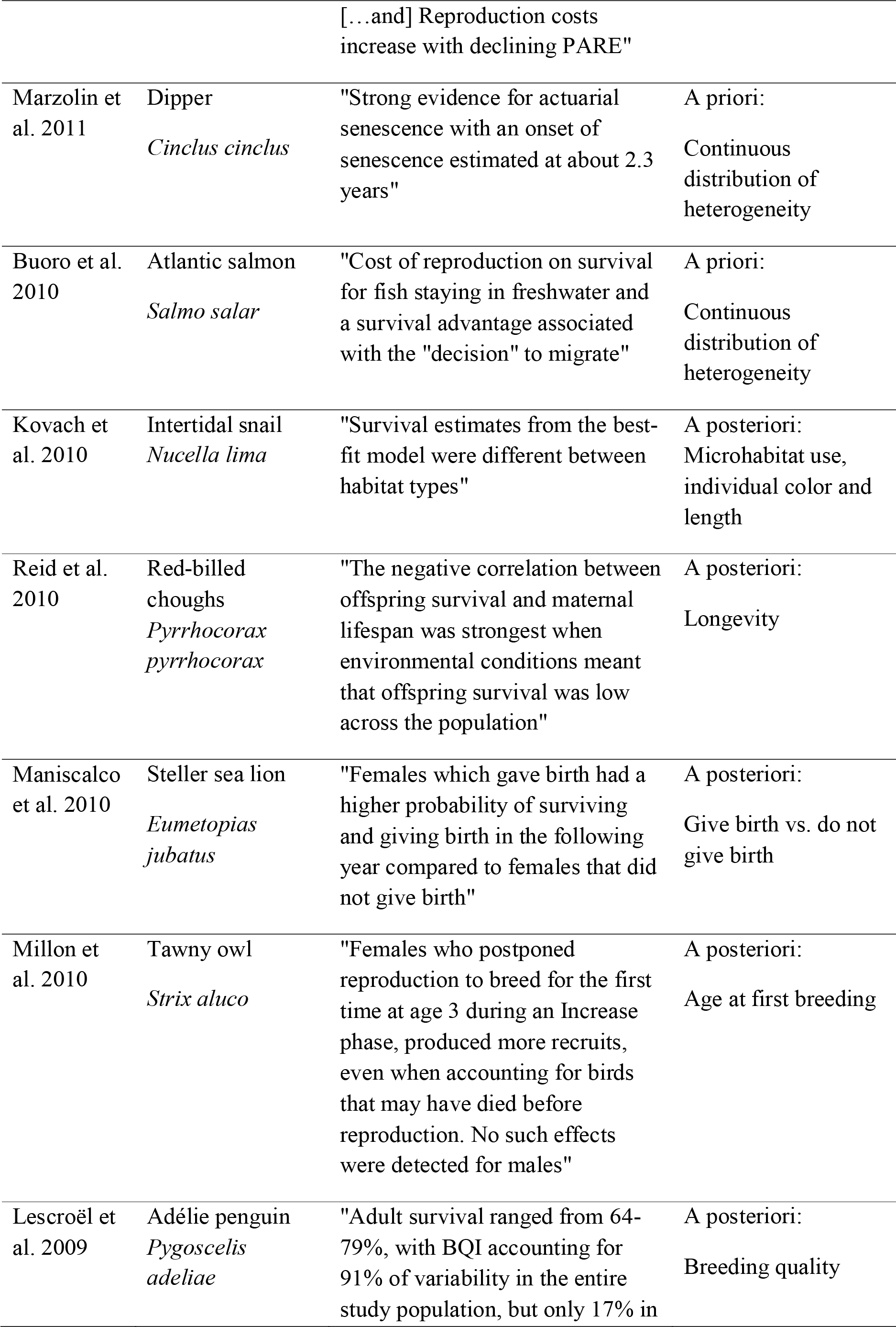
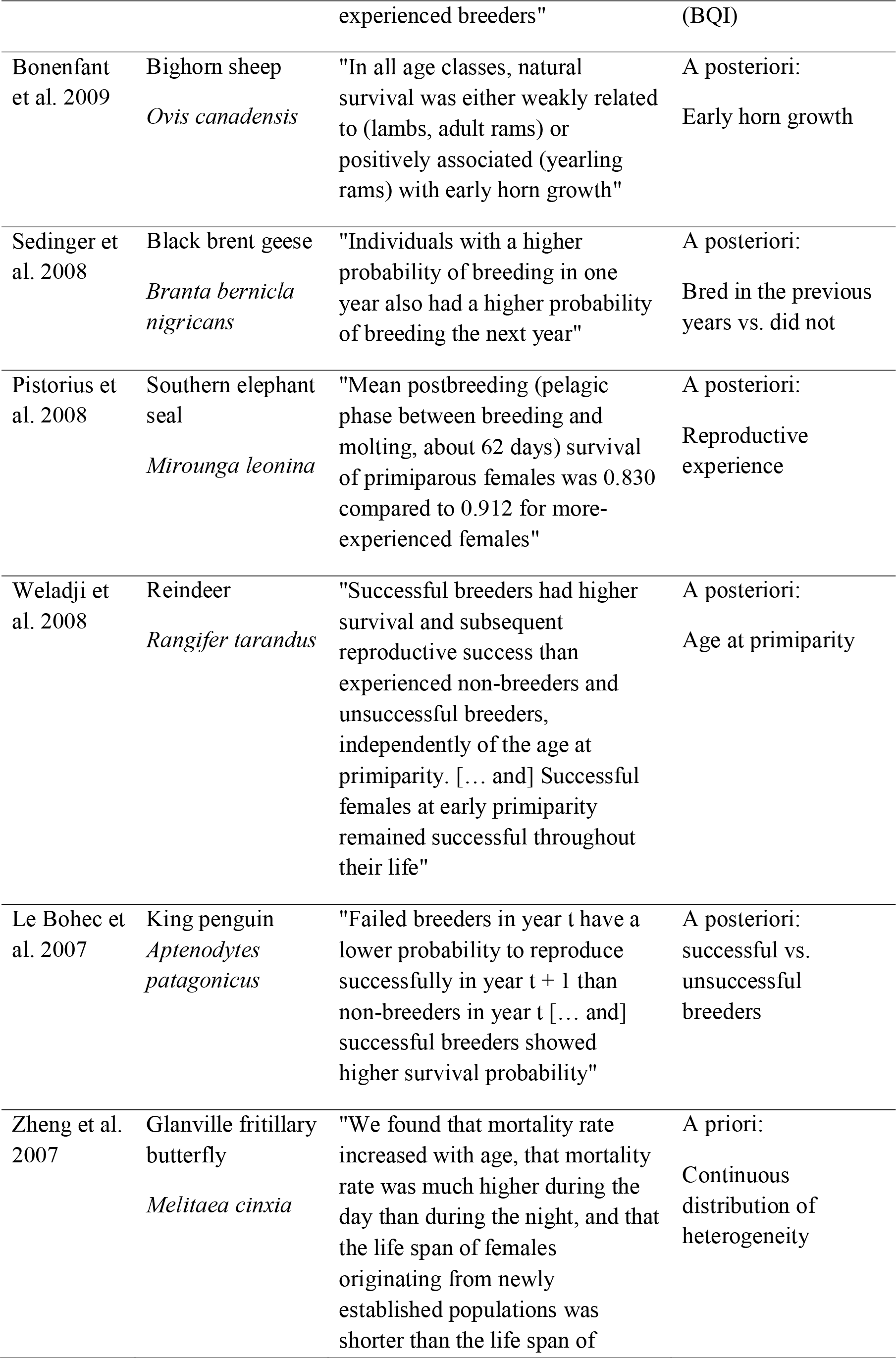
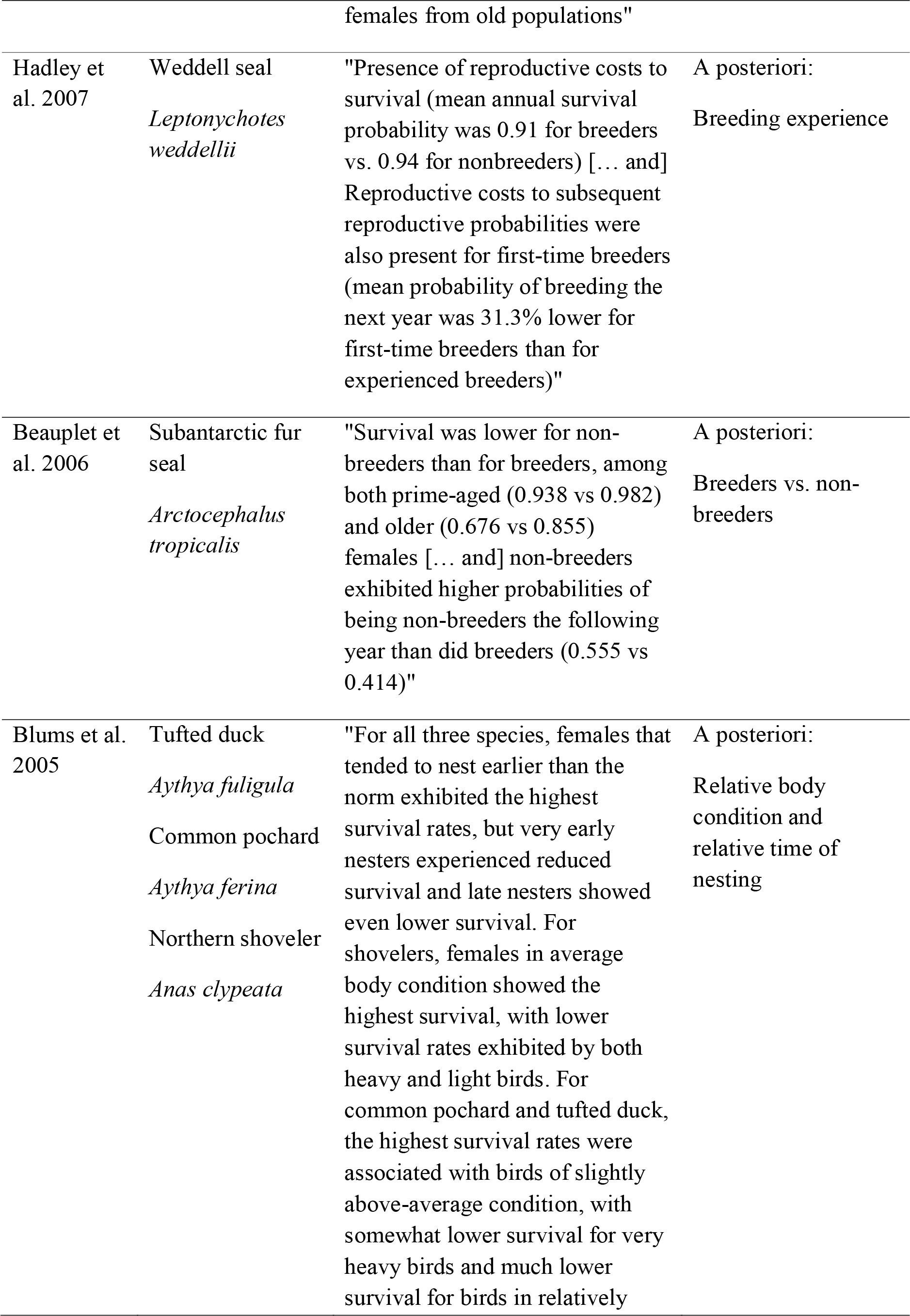
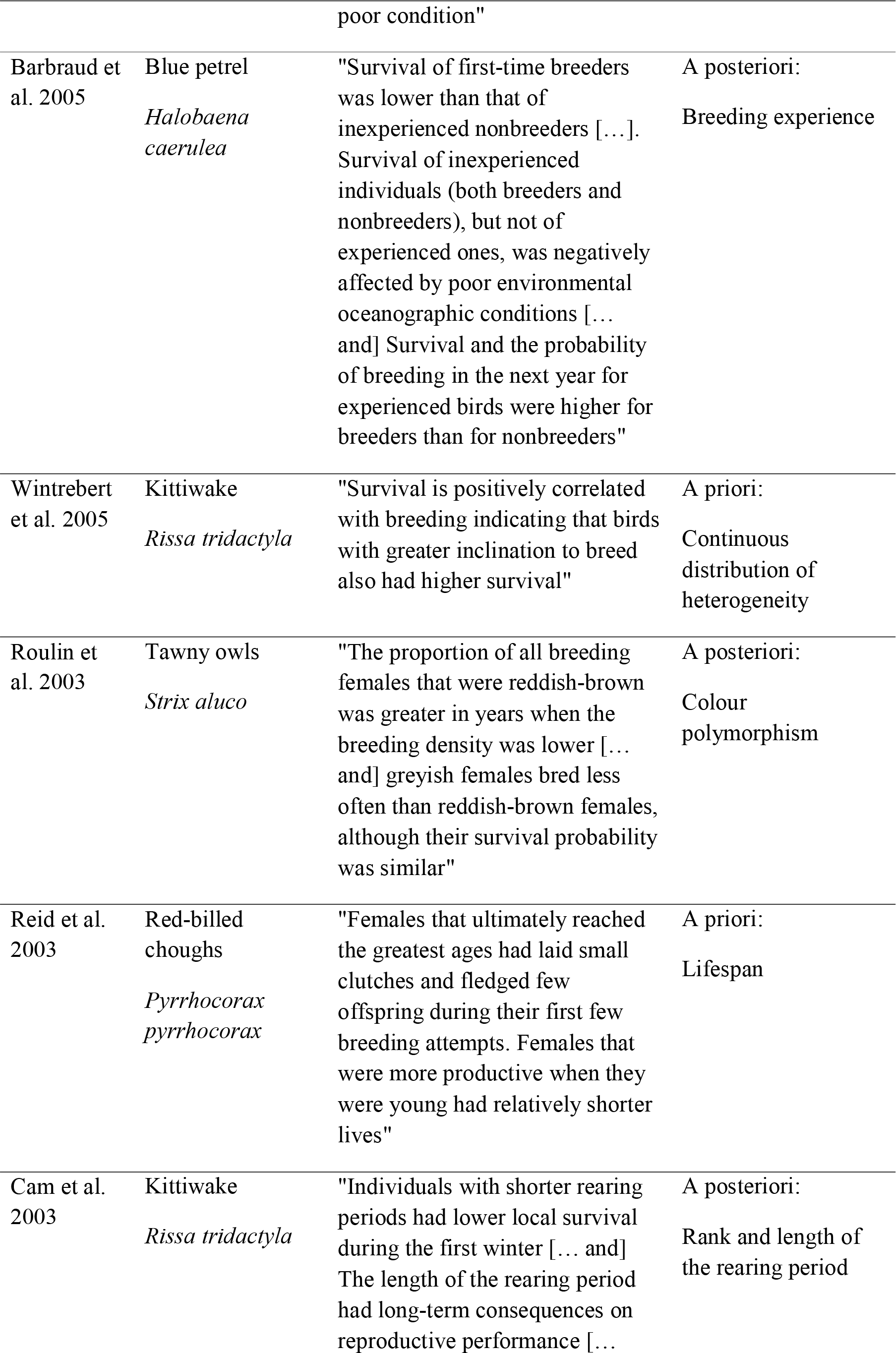
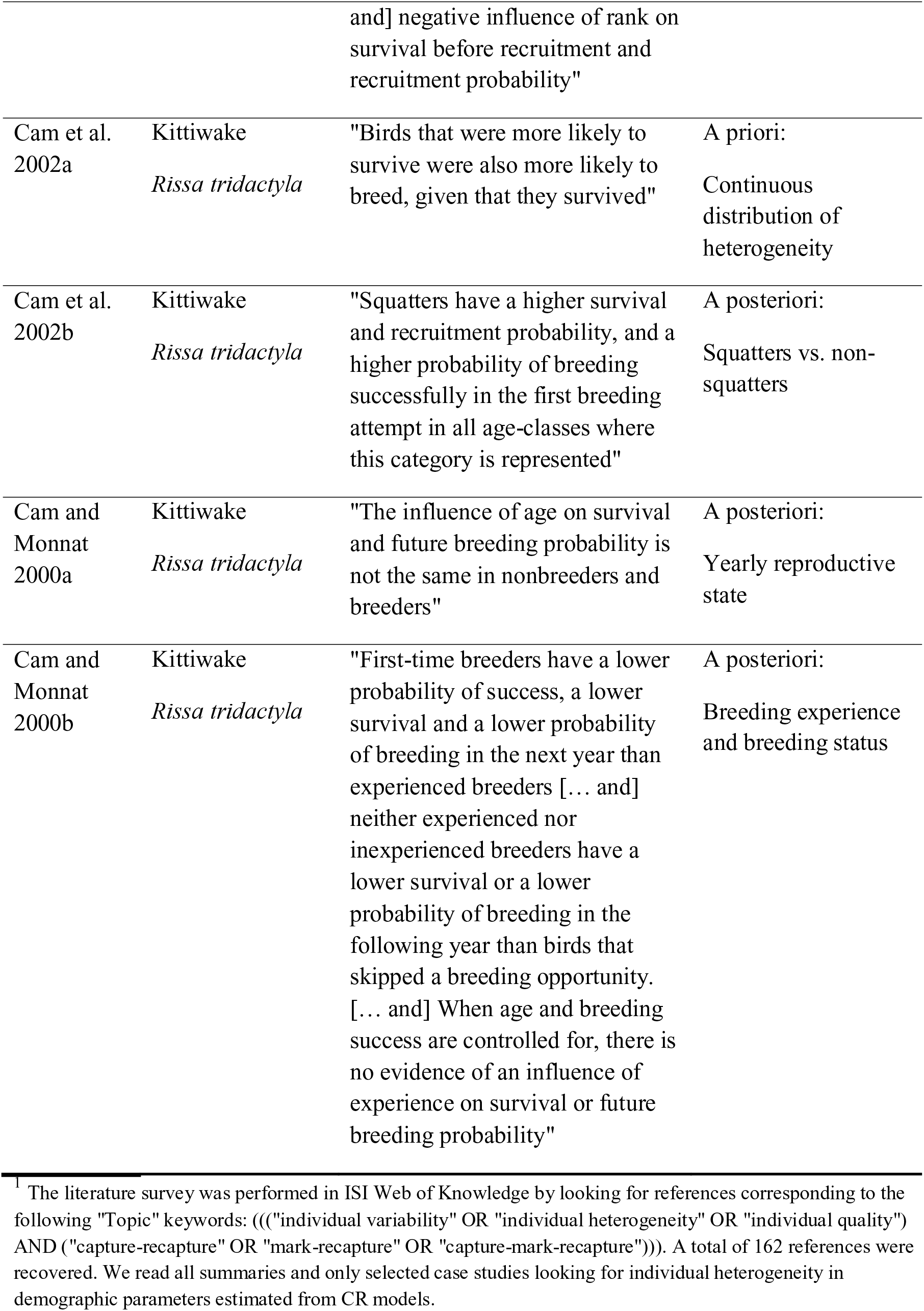
Case studies^1^ reporting an analysis of individual heterogeneity in demographic parameters (i.e. survival and reproductive traits) within a CR context. The table lists the reference, the studied species, the main outcome as explicitly stated in the paper, and how individual heterogeneity was assessed. Individual heterogeneity corresponded either to the total amount of heterogeneity (“a priori” cases) or to heterogeneity measured using some metrics (“a posteriori” cases). In these latter cases, the metrics used are provided.

We will retain Stearns (1976)’s definition of a tactic, “a set of co-adapted [demographic] traits designed, by natural selection, to solve particular ecological problems”. In the simplest case, individual heterogeneity corresponds to the random variation observed independently in each of the traits. In that case, there is no covariation between demographic traits and no life history tactic can emerge. Life history tactics can appear in response to marked differences between individuals in terms of constraints (of genetic, developmental, or environmental origin). The axis of demographic variation will thus involve a low-high continuum of performance opposing individuals weakly constrained that will perform extremely well in terms of both survival and reproduction, to individuals subjected to high constraints (sometimes called "runt", Koenig et al. 1995) that will perform extremely poorly. Individual heterogeneity in that case will lead the axis of demographic variation to correspond to a low-high continuum of individual fitness, and is often designated as a continuum of individual quality (Wilson and Nussey 2010). Alternatively, individual heterogeneity can be associated with a set of different life history tactics so that each tactic is characterized by the same mean individual fitness. For instance, some individuals will allocate a lot of energy to reproduction and pay a cost in terms of decreased survival, whereas others will allocate a lot to avoid mortality risks and pay a cost in terms of reduced reproduction, leading to the negative co-variation between survival and reproduction expected from the allocation principle (Cody 1966) to show up.

Owing to the multiplicity of factors that shape individual heterogeneity, it is impossible to account for the total amount of individual heterogeneity by measuring even a large set of traits. For a given trait, we can distinguish a *measured* individual heterogeneity using for instance phenotypic attributes such as age, sex, and size from an *unmeasured* individual heterogeneity that includes all the remaining variation for given age, sex and size (Plard et al. 2015). Until recently, this *unmeasured* individual heterogeneity was most often neglected. Assessing unmeasured individual heterogeneity is especially tricky when studying survival (or mortality) because this trait simply corresponds to a state shift for a given individual. Thus, an individual dying at 5 years of age will have survived over the first five years in a row and then will have died at 5 years, leaving the standard mixed model approach generally used for assessing individual heterogeneity in most traits (van de Pol and Verhulst 2006, van de Pol and Wright 2009) not directly applicable. However, CR models do provide a general and flexible framework for estimating and modeling both population size and demographic parameters (including survival, dispersal and recruitment) in the face of imperfect detection that is inherent to populations in the wild (Gimenez et al. 2008). These methods rely on the longitudinal monitoring of individuals that are marked (or identifiable) ideally at birth, and then encountered (*i.e*. recaptured or seen) on subsequent occasions. The first CR methods that dealt with individual heterogeneity were developed with the aim to get unbiased estimates of population size in presence of differential individual responses to trapping (Otis et al. 1978). The context has changed in recent years, and CR studies now often focus on the process of individual heterogeneity per se to assess the diversity of life history trajectories within populations, to test for the existence of different life history tactics within populations, or to assess the differential susceptibility of individuals to environmental insults.

Here we aim at providing a review of individual heterogeneity in the context of CR studies. We first define *what* we mean by individual heterogeneity by examining the landmark papers on the subject and clarifying the terminology with regards to more recent uses of the concept. Then we review the literature and illustrate *why* individual heterogeneity is used in CR studies by focusing on the detection of life-history trade-offs, including senescence. In a third section, we explain *how* to model individual heterogeneity in CR models. The distinction is made between situations in which heterogeneity can be explicitly handled by using states (e.g. breeding or disease states) or individual (time-varying or not) covariates (e.g. age or phenotype) and situations in which part of the heterogeneity remains unobserved. Regarding the latter, we outline recent developments of random-effect models and finite-mixture models. Lastly, we discuss several avenues for future research.

## What is individual heterogeneity?

### History and definitions

In CR modeling, consideration of heterogeneity between individuals of a population in demographic parameters (e.g. survival or probability of successful reproduction) has a long history: “For predictive or modeling purposes, […], heterogeneity can lead to seriously misleading conclusions, particularly if the product of two or more parameters is involved, and heterogeneity affects both of them” (Johnson et al. 1986). Early work emphasized the distinction between situations where members of populations differ with respect to some measurable attribute (e.g. sex, age), and situations where “heterogeneity is not clearly identified with a measurable variable” (Johnson et al. 1986). In the latter situation, developing methods that account for heterogeneity between individuals to estimate demographic parameters is more difficult. Early efforts toward this end echo their contemporary studies of heterogeneity in mortality risk and of aging in human demography (Vaupel et al. 1979, Manton et al. 1981, Hougaard 1984): “Unrecognized heterogeneity can lead to biased inferences, especially in time or age effects in cohort studies” (Johnson et al. 1986). In ecology, one of the earliest studies that investigated the consequences of unmeasured (and sometimes impossible to measure) variation between individuals on survival probability concerned the estimation of nest success probability using longitudinal data from nest activity (Green 1977, Johnson 1979): a nest that becomes inactive before chicks hatched is considered “dead’. The authors of early papers on CR modeling were aware of the contribution of human demographers to the development of models taking heterogeneity between individuals into account to estimate changes in mortality risk throughout life (e.g. papers cited in Johnson et al. 1986 included e.g. Keyfitz and Littman 1979, Vaupel et al. 1979 and Manton et al. 1981). According to Johnson et al. (1986), “the impact of such heterogeneity has been recognized only occasionally in animal ecology, possibly because it is difficult to deal with, and it is often relatively unimportant in many estimation problems.”

Unobserved heterogeneity can be handled using models including a continuous or discrete distribution of parameter values (recapture, breeding or detection probability). Early work by human demographers has pioneered the use of continuous distributions of mortality risks (e.g. Vaupel et al. 1979). In survival models, continuous distributions for individual heterogeneity translate the idea that individuals are characterized by a unique value of ‘mortality risk’ (or its complement, survival probability). In human demography, ‘frailty’ is traditionally used in time-to-event models, where the event of interest is death (data consist in the duration of time until death occurs). Generally, data from wild animals are longitudinal data (i.e. either they include information on whether individuals are alive or dead at each sampling occasion, or they include information on whether individuals are contacted alive at each sampling occasion, and sometimes on whether individuals are reported as dead). The idea that populations are composed of individuals that are more or less likely to experience an event (i.e., they differ in their probability of experiencing an event) is common to several areas of research. According to Wienke (2003), “Frailty corresponds to the notions of liability or susceptibility in different settings (Falconer, 1967). In the 1960’s and 1970’s, investigators developed parallel ideas in different areas of research and designed analytical methods to account for continuous distributions of “risks” in populations. In quantitative genetics, Falconer (1967) analyzed disease incidence data and assumed a continuous distribution of risks of developing the disease: “All the causes, both genetic and environmental, that make individuals more or less likely to develop a disease, can be combined in a single measure that is called ‘liability’. The liabilities of individuals in a population form a continuous variable”. In econometrics, investigators developed duration models for employment data including unobserved heterogeneity (Chamberlain 1979), where “The heterogeneity is in individual specific differences in separation rates” (i.e., the fraction of employed workers who lose jobs per time interval, see also Heckman and Borjas 1980). Heckman and Willis (1977) focused on beta-logistic models for binary data of female labor participation and assumed a random effect for unobserved individual heterogeneity in participation probability: “It is reasonable to suppose that many of the unobserved variables remain reasonably constant over time but vary considerably among women”. Obviously, the 1960’s and 1970’s stimulated the development of analytical approaches designed to handle situations where investigators acknowledge that they do not know all the relevant variables affecting individuals’ response, or where they cannot measure all of them.

Clearly, the issue raised in early CR studies is that the assumption of homogeneity of populations can lead to flawed inferences about identified parameters such as survival probability or population size (Carothers 1973). In situations where survival probability varies with age, this issue has sometimes been called a Simpson’s paradox in statistics, or an “ecological fallacy” (Kramer 1983), and has been illustrated by Cohen (1986) as follows: “The crude death rate of population A may be less than that of country B even if every age-specific death rate of country A is greater than each corresponding one of country B”. If populations are stratified according to variables that have not been considered yet, inequality of rates can be reversed, and any demographic parameter can be involved. Papers by Green (1977), Johnson (1979), Johnson et al. (1986) and the very influential paper by Vaupel and Yashin (1985) have all used analogous examples of situations with two groups to explain the consequences of unrecognized heterogeneity on inferences about mortality in ecology and human demography, respectively.

In the context of closed populations (i.e. assuming a population with no immigration, no emigration, no recruitment and no mortality), data are collected several times during the period when assumptions characterizing closed populations hold. CR models are restricted to the estimation of population size and can account for individual heterogeneity in the probability of being detected (e.g. Burnham and Overton 1979, Pollock 1980, Pollock et al. 1990, Link 2004, Pledger 2005, Farcomeni and Tardella 2010). In such a situation, “Individuals with high detection probabilities would tend to appear in the encountered sample in greater proportion than they occur in the population” (White and Cooch 2017). In CR models, the probability of detection can be assumed to vary between individuals in relation to, e.g. sex or age-related behavioral differences and more recently to space (Efford 2004, Borchers 2012). In some studies, populations are not assumed to be composed of clusters (e.g. sex, age-classes) with different detection probabilities, but each individual is “assumed to have its own unique capture probability which remains constant over all the sampling times” (Pollock 1980, Pollock et al. 1990). In CR studies where heterogeneity in the probability of detecting an individual cannot be identified using measured variables, investigators can assume that there is a distribution of individual detection probabilities and use models with individual random effects (White and Cooch 2017). In such situations, early approaches have used Jackknife estimators (Burnham and Overton 1979) or point estimators (Chao 1987). Mixture models have also been used more recently in such situations (e.g. Norris and Pollock 1996, Pledger 2000, Morgan and Ridout 2008), where populations are assumed to be composed of several hidden groups with different detection probabilities.

In the context of open populations (i.e. allowing immigration and emigration and/or recruitment and mortality to occur), models are used to estimate a variety of demographic parameters (e.g. survival rates, transition probabilities between reproductive states in successive years, populations size). Early papers (Cormack 1964, Jolly 1965, Seber 1965) have assumed homogeneous populations. Stratification according to age-classes (Manly and Parr 1968, Pollock 1981) and groups (Lebreton et al. 1992) was one of the first attempts to accommodate variation between individuals in survival probability. Arnason (1972, 1973), Schwarz et al. (1993), Hestbeck et al. (1991), and Nichols et al. (1994) laid the ground for models accounting for the fact that individuals might not belong to the same cluster during their entire life (Lebreton and Pradel 2002), but might move between states (e.g. locations, breeding states) in a stochastic manner (as opposed to movement between age-classes in a deterministic manner).

Naturally, early papers drew the same distinction as for closed populations between situations where individuals differ with respect to some measurable attribute (e.g. sex, age) and situations where “heterogeneity is not clearly identified with a measurable variable” (Pollock 1980, Johnson et al. 1986). Because open-population models can be used to estimate population size, early work also focused on the consequences of heterogeneity in detection probability on population size estimation (Carothers 1973). Just as for detection probability, two approaches have been used to account for unobserved sources of heterogeneity in survival probability in CR studies (Johnson et al. 1986): finite mixture models (Pledger et al. 2003) and random effects models (Royle 2008, Gimenez and Choquet 2010). Early studies that have assumed a continuous distribution of survival probability have also highlighted the methodological difficulties encountered in the 1990’s (Rexstad and Anderson 1992, Burnham and Rexstad 1993). In random effects models, individual heterogeneity refers to permanent differences between individuals in demographic parameters (e.g. Royle 2008, Gimenez and Choquet 2010). This definition matches exactly the concept of unobserved individual frailty proposed by Vaupel et al. (1979) where frailty designates the risk of a given individual (that is constant throughout its lifespan) to die at a given age relative to the average risk of all individuals in the population to die at this age. Up to now, most CR studies that have used mixture models for survival probability have also considered that individuals do not change cluster during their life (e.g. Fay et al. 2016), but CR models can now accommodate situations where they do (e.g. see Cubaynes et al. 2010 for an application to detection probability).

Historically dominating views of what ‘heterogeneity’ means in CR modeling depended on the class of models used. For example, ‘heterogeneity’ models in early papers focusing on closed populations referred to models where “each animal has its own unique capture probability” (Pollock 1980). Conversely, for open population models, early work on ‘heterogeneity’ considered any degree of stratification of populations, from discrete groups, or age-structured populations, to a distinct survival probability for each individual (Johnson et al. 1986). However, ‘individual heterogeneity’ has rapidly been reserved for “variation in survival probabilities among individuals after taking into account variability due to age, sex, or time” (Rexstad and Anderson 1992).

### Individual heterogeneity in contemporary CR studies

Demographic parameters are the target parameters to estimate in CR models (Lebreton et al. 1992, 2009). Other parameters such as detection probability are required to estimate demographic parameters from CR models; detection probability is relevant to sampling in wild populations, but is not a demographic parameter. However, the impact of individual heterogeneity on every type of parameter has consequences on demographic parameter estimation. CR models allow the estimation of all types of parameters.

To address individual heterogeneity in demographic parameters, the survival process (alive vs. dead), or the reproductive process (e.g. breeder vs. non-breeder or success vs. failure) is treated as a random variable. Individual heterogeneity measures differences between individuals in model parameters. Today, a broad range of approaches is used to account for individual heterogeneity in CR studies. Individual heterogeneity is understood as any source of variation between individuals in demographic parameters that cannot be accounted for by temporal or spatial heterogeneity alone, with a particular focus on the fate of individuals, their early development conditions, their ontogeny, or their past allocation to reproduction as experienced breeders. Advances in statistical methods over the past 40 years have progressively enlarged the scope of models accounting for ‘individual heterogeneity’. In CR studies, the sample scheme involves attempting to detect individuals on a discrete time basis. Contrary to continuous-time models used in human demography (Allison 1982), this specific feature must have eased the development of models in the CR arena.

Different levels of heterogeneity lead to different perceptions of ‘heterogeneity’, but the current view of ‘individual heterogeneity’ incorporates a large range of biologically relevant situations. At the lowest level individuals have their own unique demographic value (Marzolin et al. 2011; see Table 1), as in the frailty context (Vaupel et al. 1979), with a possible variation during life (e.g. Enki et al. 2014). At a broader level, heterogeneity can correspond to differences in demographic parameters between identifiable categories of individuals (e.g. identifiable groups; Drummond et al.2011), or between hidden classes or states (Péron *et al*. 2010, Pradel 2005, Johnson et al. 2016). Once included in the study, individuals can belong to the same cluster permanently (e.g. sex, hidden class; Péron *et al*. 2010) or temporarily (e.g. age, body condition, breeding state; Nichols *et al*. 1994, Pradel 2009, Johnson et al. 2016). What ‘heterogeneity’ covers in CR models inherently depends on model specification. In all cases, populations are considered as being heterogeneous.

From a biological viewpoint, individual heterogeneity in life history traits is often considered to include two components.

i. Cases in which differences between individuals are shaped early in life and are permanent during the course of the life correspond to a fixed heterogeneity. In such cases, individual heterogeneity can be accounted for by using measurable covariates a priori assumed to capture much of the individual heterogeneity (e.g. rank of offspring in birds’ clutch, Drummond et al. 2011 or body mass at the end of the maternal care period in large mammals, Hamel et al. 2009). Whether measurable individual features are translated into differences in estimates of demographic parameters, and with which method, is part of the statistical exercise. When measurable variables are missing, or insufficient to account satisfactorily for heterogeneity (Hougaard 1991), investigators can assume a discrete or continuous distribution of demographic parameters (Royle 2008, Péron et al. 2010). In agreement with the concept of frailty, investigators assume that there are differences in demographic parameters between individuals that cannot be associated with measurable covariates and use latent variables to quantify them (e.g. Hougaard 1995, Yashin et al. 2008, Cam et al. 2013, Hamel et al. 2014, Cam et al. 2016).
ii. However, not all individual heterogeneity is fixed. Individual differences in a given trait at a given time are subjected not only to the influence of early-life conditions, but also to current conditions both at that time and between early life and that time. As above, in some situations, observable variables are available to account for variation in individuals’ demographic parameters throughout their life (e.g. age: Pollock 1981; group: Lebreton *et al*. 1992; state: Nichols *et al*. 1994). However, when such observable variables are missing or inefficient at capturing most individual heterogeneity (Hougaard 1991), unobserved, latent traits changing over life can be used. Such cases correspond to ‘dynamic frailty’ (Pennell and Dunson 2006, Duffie et al. 2009, Chambert et al. 2013, Cam et al. 2004).

Irrespective of the component shaping individual heterogeneity in a given population, the total amount of individual heterogeneity in that population is not constant and varies over time. Thus, Lomnicki (1978) pointed out that asymmetric responses of individuals to increased competition that occur in presence of harsh environmental conditions (Lomnicki developed his argument in the context of density-dependence but the same pattern is expected whatever the cause of resource limitation) lead individual heterogeneity within a population to increase.

## Why individual heterogeneity in a CR context?

### Individual heterogeneity seen as a nuisance

Because analyzing most CR datasets requires using models that include detection probability, the existence of individual heterogeneity in this parameter has stimulated a large number of works in the early CR literature (e.g. Otis et al. 1978). As Eberhardt (1969) pointed out, “various sets of data indicate that the equal-probability-of capture assumption is not fulfilled.” Unequal detection rates may lead to biases in abundance estimate, estimated survival probability or population growth in the context of open populations (e.g. Carothers 1973, Schwarz 2001). Heterogeneity in detection probability is still receiving attention in contemporary studies (Crespin et al. 2008, Cubaynes et al. 2010, Pradel et al. 2010, Marescot et al. 2011, Oliver et al. 2011, Fletcher et al. 2012, Abadi et al. 2013, White and Cooch 2016), and new methods are being developed to address heterogeneity, such as multievent models (Pradel 2009).

### Individual heterogeneity as a biological process

Methodological developments of CR models have considerably increased the relevance of CR studies to address questions not only in ecology, but also in evolution. A key feature of this development is the increased ability of CR models to account for variation between individuals in demographic parameters as well as detection probability at the scale assumed to be relevant in the studied context. This scale can be the individual, rather than permanent groups (e.g. sex), or temporary aggregates of individuals (e.g. reproductive states). The development of evolutionary biology as a powerful conceptual and methodological framework for biological disciplines (e.g. Dobzhansky 1973) has brought a new perspective on individual heterogeneity. In CR studies, instead of being addressed because of potential biases in estimates of abundance, survival probability or population growth rate (e.g. Crespin et al. 2008, Cubaynes et al. 2010, Pradel et al. 2010, Oliver et al. 2011, Abadi et al. 2013), individual heterogeneity has become the focus of studies because of its biological relevance. For evolutionary biologists, the individual level can be relevant to address natural selection if heritable variation is expressed at this level (Chambert et al. 2014). In addition, variation between individuals in demographic parameters is relevant to population ecology and dynamics, whether it concerns traits that are heritable, or not, and whether it can be accounted for using observed variables, or not (e.g. Kendall and Fox 2002). Indeed, both population extinction risk and viability depend on the degree and structure of individual heterogeneity in survival probability and reproductive parameters (Conner and White 1999, Stover et al. 2012).

Starting from classes of models where demographic parameters varied with time (Jolly 1965), groups of individuals, or age, the development of software programs to build multistate models (Arnason 1972, 1973) in the 1990’s has considerably increased the attractiveness of CR models. These models indeed allow biologists to address questions about a large range of factors structuring a population, which determine individual sequences of states between which individuals move in a stochastic manner (Nichols et al. 1994, Nichols and Kendall 1995). These models triggered studies of life histories using CR models (e.g. Cam et al. 1998, Hadley et al. 2007). Another class of approaches, multievent models (Pradel 2005), has also helped biologists address questions about the influence of ‘state’ on demographic parameters (Sanz-Aguilar et al. 2011). Indeed, one of the difficulties in CR studies is that ‘state’ may not be observed with certainty, or even not be observed at all (Desprez et al. 2013). Moreover, the development of user-friendly software to build models with individual covariates has also stimulated work in evolutionary ecology using CR data (e.g. Gimenez et al. 2009). The question of time-specific individual covariates with missing values is still a current issue (Bonner and Schwarz 2006). The ability of CR models to accommodate variation between individuals in demographic parameters raises the issue of methods of inference about model parameters in CR studies (e.g. Pledger and Schwarz 2002, Royle 2008, Gimenez and Choquet 2010). This issue is tightly linked to the level of stratification of populations, or of aggregation of observations (Cooch et al. 2002). As emphasized by Nichols (2002): “If we view an individual organism’s fate or behavior at any point in space and time as a unique event not capable of informing us about the likelihood of the event for other individuals or points in space and time, then generalization and prediction become impossible”. To allow formal statistical inferences about variation in demographic parameters and detection probability in populations, CR models rely on assumptions, notably regarding unobserved heterogeneity (e.g. a distribution of random effects; Royle 2008, or a mixture model; Péron et al. 2010, Marescot et al. 2011).

### Assessing senescence in the wild: an increasingly popular focus of CR studies

The process of senescence, which can be interpreted in the context of the allocation principle as the trade-off opposing performance during early life and performance in late life (Baudisch and Vaupel 2012, Lemaître et al. 2015), has been the focus of a large number of empirical studies during the last decade (see Nussey et al. 2013 for a review). As imperfect detection of individuals is the rule in free-ranging populations (Gimenez et al. 2008), CR has become the gold standard to measure reliably actuarial senescence in the wild (e.g. Loison et al. 1999, Bouwhuis et al. 2012). The question of level of inference has recently emerged as a critical point in CR studies of senescence (Péron et al. 2010, Marzolin et al. 2011). Based on early work by human demographers addressing heterogeneity in mortality risk (e.g. Vaupel et al. 1979, ecologists have often used the concept of frailty. However, Vaupel and Yashin (1985) considered the case of a heterogeneous population with two classes of individuals, frail and robust ones. As time passes and individuals age, there is a disjunction between the variation of the mean survival probability (i.e., when pooling frail and robust individuals) with age, and the variation in survival probability with age within each group (Figures 1, 2). Ignoring heterogeneity in mortality risk may lead to flawed inferences about aging rate (Vaupel and Yashin 1985, Zens and Peart 2003), a phenomenon documented in wild animals using CR models (Nussey et al. 2008, Péron et al. 2016). This phenomenon has long been acknowledged in wildlife studies for life stages other than senescence (e.g. nest mortality; Green 1977 and Johnson et al.1986, and Burnham and Rexstad 1993 in the context of CR studies). The consequences of ignoring individual heterogeneity in survival probability have also been investigated in CR studies using special datasets with perfect detection of individuals (e.g. Cam et al. 2002a, 2013, Wintrebert et al. 2005, Fox et al. 2006, Aubry et al. 2011, Knape et al. 2011).

The requirement of accounting for heterogeneity in survival studies was raised by human demographers very early (which distribution to use to account for individual heterogeneity in mortality risk, Manton et al. 1986) and has become a key topic in ecology. Some demographers argued that observable criteria might not account for individual heterogeneity in a satisfactory manner, and developed mixed models or mixture models for time to event data (Kannisto 1991, Abbring and Van Den Berg 2007). The debate about the appropriate distribution to consider is also taking place in ecology (e.g. Gimenez et al. 2010, Péron et al. 2010, 2016). However, in CR studies, biologists use discrete data (e.g. mixed binomial models for survival), which may lead to fewer issues with parameter identifiability and assumptions than hazard models with frailty (Wienke 2010). Moreover, to some extent, the idea of addressing heterogeneity using a distribution of latent demographic traits is coherent with approaches to quantify variation in populations that are familiar to evolutionary biologists, namely, variances in traits in quantitative genetics (Lynch and Walsh 1998, Chambert et al. 2013, 2014). Recently capture-recapture animal models (CRAM) have been developed to estimate heritability of demographic parameters (Papaïx et al. 2010).

### Detection of trade-offs between life history traits

Trade-offs are one of the cornerstones of the theory of life histories (Roff 1992). They are based on the principle of allocation (Cody 1966) and express the idea that individuals possess a limited amount of energy and have thereby to share energy among various functions so that individuals allocating a lot of energy into current reproduction cannot allocate as much into survival or future reproduction (Roff 1992). However, empirical analyses have often failed to detect trade-offs in the wild because of individual heterogeneity in resource acquisition. van Noordwijk & de Jong (1986) indeed demonstrated that positive associations between current reproduction and future survival or reproduction occur when individual heterogeneity in resource acquisition is greater than individual heterogeneity in resource allocation. The development of multistate models has attracted evolutionary biologists to study trade-offs within the CR arena (Cam et al. 1998, Yoccoz et al. 2002). Individuals are assumed to make allocation decisions according to their own state (McNamara and Houston 1996). Consequently, any unobserved feature of ‘state’ may explain why trade-offs are not detected. Experimental approaches may help unveil trade-offs (Reznick 1985), but may also go against heterogeneity (e.g. Festa-Bianchet et al. 1998, Yoccoz et al. 2002). At the extreme, each individual can be assumed to be in a unique ‘state’ that cannot be measured, and trade-offs might not be detected. Some observational CR studies have provided evidence of trade-offs after identifying traits that reliably described changes in individual state (e.g. social rank, mass, etc., Hamel et al. 2009), by taking advantage of unfavorable conditions (e.g. Descamps et al. 2009) or by distinguishing direct from indirect effects (Cubaynes et al. 2012a). The development of hierarchical CR models with individual heterogeneity has allowed investigators to assume a distribution of latent life history traits in populations (Royle 2008, Gimenez and Choquet 2010). In particular, Buoro et al. (2010, 2011) have been successful at detecting trade-offs using this type of approach.

## How to infer individual heterogeneity in CR models

In this section, we provide details about the CR models used in the case studies reviewed above. Specifically, we focus on multistate, random-effect and finite-mixture CR models possibly including individual covariates because these are currently the most commonly used tools to incorporate individual heterogeneity and deal with detectability less than one. We focus on survival and open populations in the following tutorial, but the methods are applicable to other CR model parameters (e.g. Matechou et al. 2016), including the detection probability, and in other contexts such as closed populations. For the sake of illustration, we simulate data in R that we analyze i) in a frequentist framework using maximum likelihood methods with program MARK (White and Burnham 1999) called from R using the package RMark (Laake 2013; alternatively, see the R package marked by Laake et al. 2013) and E-SURGE (Choquet et al. 2009) and ii) in a Bayesian framework using Markov chain Monte Carlo methods with program JAGS (Plummer 2003) called from R using the package R2Jags. Below we present results from the frequentist approach only. The code to simulate data and fit CR models is available in the Supplementary materials and from GitHub (https://github.com/oliviergimenez/indhet_in_CRmodels).

### *Measured* individual heterogeneity: individual covariates and multistate CR models

#### Individual covariates

We start with a simple example of an individual *i* with the encounter history *h_i_* = 101 where ‘1’ is for detected and ‘0’ for non-detected. Here, individual *i* was detected on the first sampling occasion, then missed and eventually detected again on the last sampling occasion. We consider the Cormack-Jolly-Seber model for open populations and assume that neither survival probability *ϕ between* two sampling occasions nor detection probability *p at* a sampling occasion vary between individuals. Then, the contribution of individual *i* to the model likelihood is:

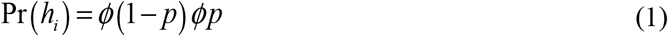

Now let us assume that we are able to measure individual heterogeneity under the form of an individual covariate, say *x_i_*, which takes a specific value for individual *i* (Pollock 2002). We assume the covariate to characterize the individual throughout the CR study (i.e., it is not a time-varying covariate, see below). Then, individual variation in the survival probability (or the detection probability) can be partly explained by this covariate through:

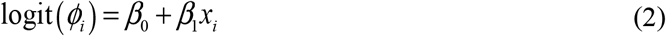

where *ϕ_i_* is the survival probability for individual <u>*i*</u>, 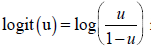 is the logit function and is used here as a constraint to make sure that survival is estimated between 0 and 1, and the *β*’s are regression coefficients to be estimated (e.g. North & Morgan 1979). Assuming now a model with individual-specific survival, Eqn (1) becomes:

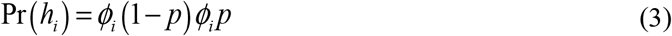

We do not estimate survival for each individual, but instead the regression coefficients *β*’s in Eqn (2) by first using the reciprocal logit function 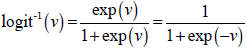 in Eqn (2) and plugging in the result in Eqn (3):

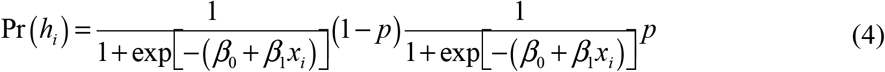

The covariate *x_i_* may be continuous such as body mass or discrete such as sex. If the covariate is discrete, it is usually referred to as a *group* in the CR literature (Lebreton et al. 1992).

So far, we have assumed that this covariate does not vary over time, in other words that an individual *i* has the same value *x_i_* of the covariate whatever the sampling occasion (i.e. matching with the concept of frailty *sensu stricto*). When dealing with time-varying individual covariates, which matches the concept of dynamic frailty (for unobserved heterogeneity), we then need to distinguish between discrete and continuous covariates.

#### Discrete time-varying individual covariates and multistate CR models

Discrete time-varying individual covariates are referred to as *states* in the CR literature (e.g. breeder/non-breeder or infected/non-infected), and are analyzed with so-called multistate CR models (Schwarz et al. 1993, Lebreton et al. 2009). Let us assume that we measure a time-varying individual covariate with two levels, A and B, and that individual *i* has now the encounter history *h_i_* = A0B with obvious interpretation. Two things might have happened on the second sampling occasion at which the individual was not detected: either it stayed in state A or it made a transition to state B. The transition event immediately calls for the introduction of additional parameters, namely the transition probability *ψ^AB^* from state A to state B and *ψ^BA^* from state B to state A. The probability of staying in state A (or B) is obtained as the complementary probability *ψ^AA^* = 1 *− ψ^AB^* (or *ψ^BB^* = 1 −*ψ^BA^*). The two events ‘being alive in state A’ and ‘being alive in state B’ at the second sampling occasion cannot occur together: these are mutually exclusive. As a result, the contribution of individual *i* to the model likelihood has two components depending on the actual underlying encounter history AAB or ABB:

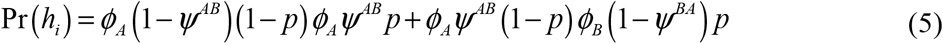

Note that *p* does not depend on state for simplicity, but this does not need to be the case. An example of the use of multistate CR models to detect life-history trade-offs in the presence of individual heterogeneity is provided in Table 2.

**Table 2.** Detection of a trade-off between reproduction and survival using multistate capture-recapture models after individual heterogeneity is accounted for. We simulated multistate capture-recapture data with two states, non-breeding (NB) and breeding (B). To mimic individual heterogeneity, we considered robust individuals with survival 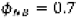 and 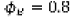 and frail individuals with survival 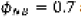 and 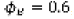, the only difference being in the survival of frail breeders that is much lower than that of robust breeders. For each group, we simulated the fate of 100 newly marked individuals in each year of a 6-year experiment. We report parameter estimates from two multistate models in which i) we ignored individual heterogeneity (column ‘*Ignoring* individual heterogeneity) and ii) we explicitly incorporated an individual covariate to handle this source of heterogeneity (column ‘*Incorporating* individual heterogeneity’). The parameters we used to simulate the data are given in the column ‘Truth’. We refer to the Appendix for more details. **The cost of breeding on survival is detected only in frail individuals after accounting for individual heterogeneity through quality** 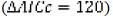.

| Parameter | *Ignoring*<br>Individual heterogeneity | *Incorporating*<br>individual heterogeneity | Truth |
| --- | --- | --- | --- |
| Survival of frail non-breeders | 0.69 [0.67; 0.72] | 0.70 [0.66; 0.73] | 0.7 |
| Survival of frail breeders | 0.70 [0.68; 0.72] | 0.58 [0.55; 0.61] | 0.6 |
| Survival of robust non-breeders | 0.69 [0.67; 0.72] | 0.69 [0.65; 0.72] | 0.7 |
| Survival of robust breeders | 0.70 [0.68; 0.72] | 0.80 [0.77; 0.82] | 0.8 |
| Transition from non-breeding to breeding | 0.78 [0.75; 0.80] | 0.78 [0.75; 0.80] | 0.8 |
| Transition from breeding to non-breeding | 0.31 [0.29; 0.33] | 0.31 [0.28; 0.33] | 0.3 |
| Detection | 0.90 [0.88; 0.91] | 0.90 [0.88; 0.91] | 0.9 |

#### Continuous time-varying individual covariates

Continuous time-varying individual covariates are difficult to deal with. Ideally, we have:

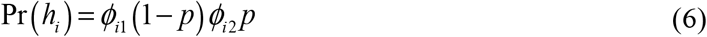

with

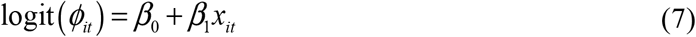

where *ϕ_it_* is the survival probability for individual *i* between sampling occasions *t* and *t* + 1 and *x_it_* is the value of the covariate for individual *i* at occasion *t*. However, this only corresponds to the ‘ideal’ situation because when an individual is not detected at a particular sampling occasion, then the value of the covariate is generally unknown, which makes it impossible to form Eqn (7). A first possibility is to omit individuals with missing values or to replace the missing values by, for example, the mean of all covariate values observed for an individual. These ad-hoc approaches result in a loss of information and bias in parameter estimates and should be avoided (Kendall et al. 2003, Lee et al. 2016). A more formal approach consists in imputing missing covariate values from an underlying distribution that is used to model the change in covariate values over time, typically a first-order Markov process such as a random walk (Bonner & Schwarz 2006, Langrock & King 2013; see also Worthington, King & Buckland 2015). A second possibility involves the discretization of the covariate in two or more levels so that multistate CR models can be used (Nichols et al. 1992). Lastly, inference can be based on a conditional likelihood approach using only the observed covariate values – the so-called trinomial approach (Catchpole et al. 2010). Several studies have compared the statistical performances of these methods (Bonner et al. 2010, Langrock and King 2013) and found that imputation methods were sensitive to the covariate model and that all methods were sensitive to the detection probability and the number of missing values. In practice, discretizing the continuous covariate and using multistate CR models is a pragmatic approach that can easily be implemented in existing software packages.

### *Unmeasured* individual heterogeneity: random-effect and finite-mixture CR models

If for some reason, heterogeneity cannot be measured, or there is a reason to believe that individual covariates do not capture the relevant variation, it can yet be incorporated using two approaches.

#### Random-effect CR models

The usual random-effect approach has been adapted to CR models (Coull and Agresti 1999, Royle 2008, Gimenez and Choquet 2010). We write:

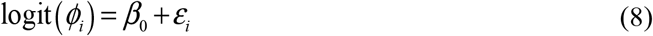

where the *ε_i_*’s are normally distributed with mean 0 and variance *σ*^2^ to be estimated, which is to be plugged in Eqn (3) using the reciprocal logit function. To fit this random-effect model, one can adopt a Bayesian (Royle 2008) or a Frequentist approach (Gimenez and Choquet). More complex structures in the random effects can be considered (heritability: Papaïx et al. 2010a; nested effects: Choquet et al. 2013). An example of the use of random-effect CR models to detect senescence in the presence of individual heterogeneity is provided in Figure 1.

**Figure 1.**
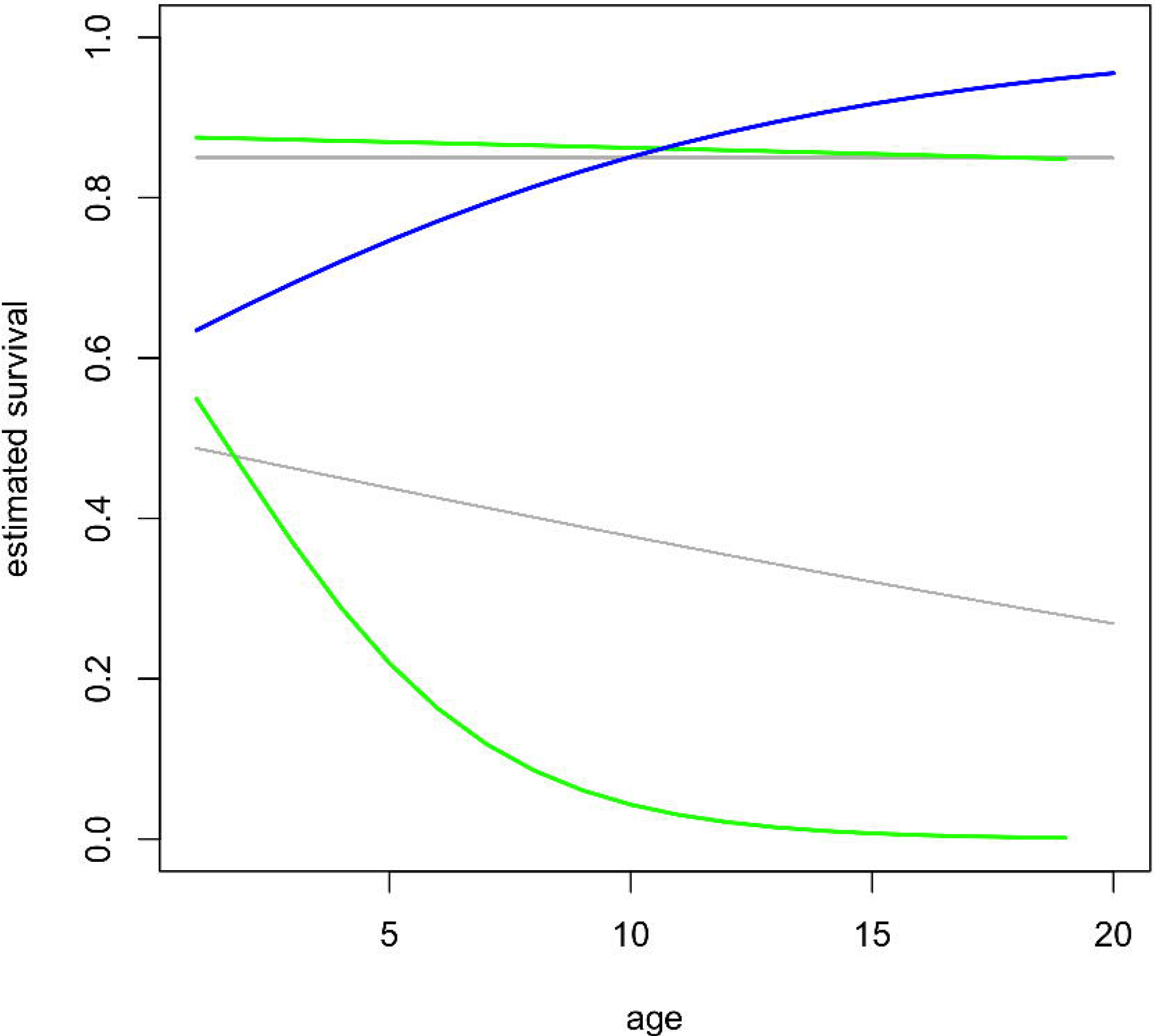
Senescence is masked when individual heterogeneity is not accounted for: random-effect capture-recapture model. We simulated the fate of 500 individuals (in grey) from a single cohort with survival decreasing as they age over a 20-year study. We also added a frailty for each individual 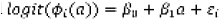 where 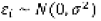. We used 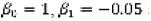 and σ = 1. We considered the same detection probability *P* = 0.5 for all individuals. We report the age-specific survival patterns from two models in which i) we ignored individual heterogeneity (in blue) and ii) we incorporated an individual random effect to handle with this source of heterogeneity (in green), both to be compared to the actual trend that we used to simulate the data (in red). Clearly, ignoring individual heterogeneity obscures senescence in survival. We refer to the Appendix for more details.

#### Finite-mixture CR models

Another avenue to handle with unobserved individual heterogeneity is to use finite-mixture models (Pledger et al. 2003, 2010, Pledger 2005, Pledger and Phillpot 2008). These models assume that individuals can be categorized into a finite number of heterogeneity classes (hidden states). More explicitly, an individual may be alive in class C_1_ or class C_2_. Then,

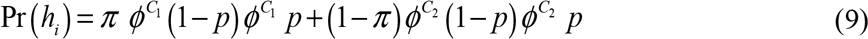

where *π* (resp. 1 − *π*) denotes the proportion of newly marked individuals in class C_1_ (resp. C_2_). Transition between classes can be considered (Pradel 2009; see below). An example of the use of finite-mixture CR models to detect senescence in the presence of individual heterogeneity is provided in Figure 2.

**Figure 2.**
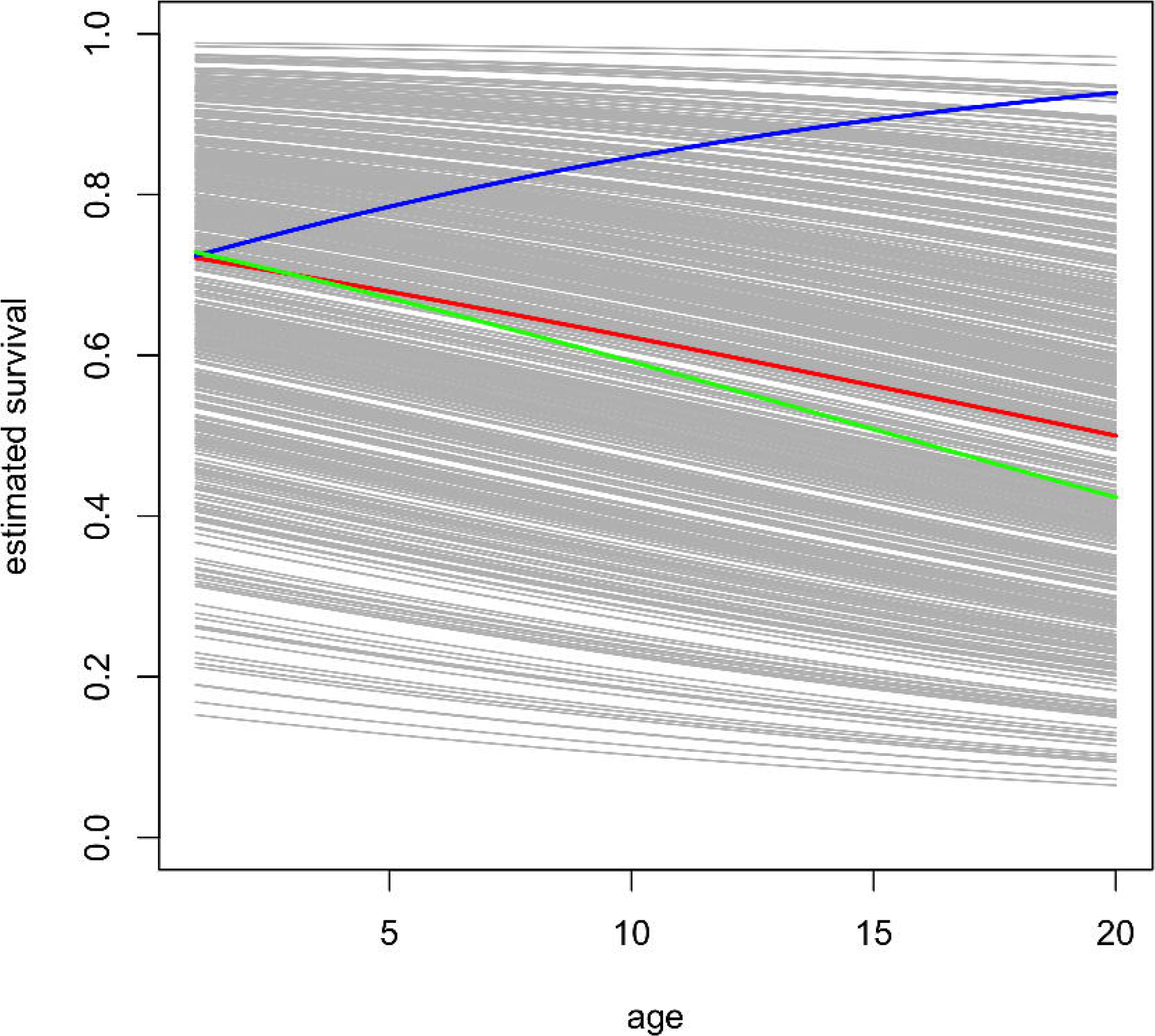
Senescence is masked when individual heterogeneity is not accounted for: finite-mixture capture-recapture model. We simulated the fate of 1000 individuals from a single cohort that were split into a group of robust individuals in proportion π with constant high survival 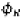 and a group of frail individuals with survival 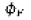 that aged over the 20 years of the study according to the relationship 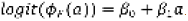. We used π = 0.5, 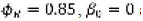 and 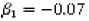. We considered the same detection probability 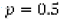 for all individuals. We report the age-specific survival patterns from two models in which i) we ignored individual heterogeneity (in blue) and ii) we used mixture models with two hidden classes of individuals to handle with heterogeneity (in green), both to be compared to the actual trend that we used to simulate the data (in grey). Clearly, ignoring individual heterogeneity obscures senescence in survival. We refer to the Appendix for more details.

### Hidden-Markov modeling framework

CR models can be fruitfully expressed as state-space models in which the biological process (survival for example) is explicitly distinguished from the observation process (detection) (Gimenez et al. 2007, 2012, Royle 2008, King 2012). In particular, multistate CR models incorporating uncertainty in state assignment – multievent CR models – have been formulated as hidden-Markov models (HMM; Zucchini *et al*. 2016) by Pradel (2005; review in Gimenez *et al*. 2012), a particular case of state-space models in which the states are Markovian (i.e. the next state depends only on the current state and not on the sequence of states that occurred before). An advantage of the HMM formulation of CR models is that it provides high flexibility in the way individual heterogeneity is modeled. For example, the HMM formulation of finite-mixture CR models can easily be extended to consider transitions between classes of heterogeneity (Pradel 2009, Cubaynes et al. 2010). Let us define the states alive in class 1 (‘C_1_’), alive in class 2 (‘C_2_’) and dead (‘D’). The individuals can go undetected (‘0’) or detected (‘1’). Initially, the state of an individual is driven by the vector of initial state probabilities:

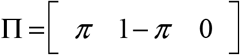

where the states C_1_, C_2_ and D are in columns in that order. Then the observation process at first capture applies through:

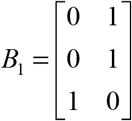

where the states are in rows and the observations (or events) are in columns (0 and 1 in that order). Now that the fate of individuals at first capture occasions is modeled, the survival and observation processes occur successively at the subsequent occasions. The survival process is governed by:

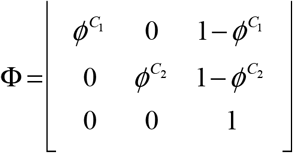

where the states at *t* are in rows and the states in *t* + 1 are in columns. Individuals can be allowed to move from one heterogeneity class to the other through transition probabilities *ψ* by multiplying the survival matrix by a transition matrix:

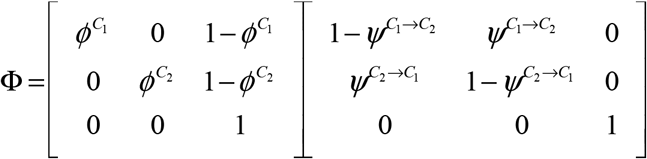

The observation process at occasion *t* is modeled using:

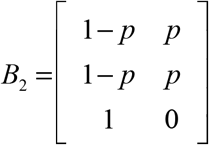

where states are in rows and observations in columns. The probability in Eqn (9) can be written as the product of the matrices above (Pradel 2005).

### Is individual heterogeneity statistically relevant?

Testing the statistical relevance of individual heterogeneity can be done in two ways. First, the quality of fit of models with heterogeneity to CR data can be assessed using goodness-of-fit tests. An ad-hoc procedure was proposed in the context of finite-mixture models by considering specific combinations of components of the goodness-of-fit test for homogeneous models (Péron et al. 2010). A more formal approach is being developed (Jeyam, McCrea, Pradel, unpublished results) based on methods used in behavioral sciences. Second, models with and without heterogeneity can be compared using hypothesis testing or model selection. For multistate models, model selection using the Akaike Information Criterion (AIC; Burnham & Anderson 2002) is usually favored as illustrated in Table 2. In the Bayesian context, several methods have been used and we refer to review papers for guidelines (O’Hara and Sillanpää 2009, Tenan et al. 2014, Hooten et al. 2015). In random-effect models, the question boils down to testing whether the variance of the random effect is zero, which can be addressed using likelihood ratio tests (Gimenez and Choquet 2010) but may be difficult to do in a model selection framework (Bolker et al. 2009). We refer to O’Hara & Sillanpää (2009) for Bayesian methods (see also Royle 2008, Chambert et al. 2014). In finite-mixture models, standard tools from the model selection framework, namely the AIC, can be used (Cubaynes et al. 2012b), although it may fail in the context of detecting senescence (Supplementary Material A in Péron et al. 2016). In a Bayesian context, the Deviance Information Criterion (DIC; Spiegelhalter et al. 2014) is known to perform poorly on mixture models, and the Watanabe-Akaike information criteria (wAIC; Gelman et al. 2014) holds promise in this context, although it is yet to be used with CR models.

## Discussion and research perspectives

### Random effects vs. mixture?

In studies based on CR models built in the context of closed or open population CR models, as well as in human demography, there is currently a debate about the distribution to consider to account for individual heterogeneity in demographic parameters (Yashin et al. 2001, Péron et al. 2010). This debate sometimes focuses on the biological justification of continuous distributions *vs*. mixture models (e.g. Péron et al. 2010). The debate also focuses on alternatives to distributions that might be unrealistic or inadequate (Péron et al. 2010). This question is not specific to CR modeling (Hamel et al. 2016). Clusters of individuals sharing values of latent traits can be identified using mixture models. Recently, Hamel et al. (2016) addressed the question of the identification of reproductive and growth tactics in long-lived mammals using mixture models. They also used simulations and showed that in many cases the number of clusters can be chosen using an information theoretic or a bootstrap approach. Alternatively, infinite mixture models could be developed for CR data (Rasmussen 2000, Ohlssen et al. 2007, Raman et al. 2010), where the number of clusters is *a priori* very large, but the number of clusters including at least one individual is estimated and can range from 1 to a large number; the latter situation leads to a distribution of demographic parameters that approaches a continuous one (Ohlssen et al. 2007). Rather than comparing models that vary in complexity using for instance an information criterion, Bayesian nonparametric approaches fit a single model that can adapt its complexity to the data (Gershman & Blei 2012; see Ford, Patterson & Bravington 2015 and Manrique-Vallier 2016 for applications to CR models). Moreover, the question of how to account for heterogeneous detection probability in CR models designed to estimate population size has a very long history (e.g. Carothers 1973, Link 2003, Ghosh and Norris 2005). Carothers (1973) investigated the consequences of violations of the assumption of equal detection probability on estimates from the Jolly-Seber model. He concluded that “any distribution is, from the point of view of investigating bias, as good as any other with the same [mean detection probability] and [coefficient of variation], and it is therefore justifiable to select a distribution on the grounds of computational convenience alone”. The number of classes might be itself of interest, but in the framework of closed populations, there is no straightforward means of determining the number of components of a mixture model for detection probability (Link 2003), and it is strongly advised against trying to interpret the mixture parameters (Shirley Pledger, pers. comm.).

### Change in latent values of demographic parameters over lifetime

In standard models for longitudinal data with individual heterogeneity, an independent subject-specific random effect is assumed to be constant over time for each subject (Vaupel & Missov 2014), which matches early versions of the concept of frailty (Vaupel et al. 1979). Generally, in CR studies using mixture models, each individual is also assumed to be a member of a latent class when it enters the study, and it does not change class. Mortality risk or breeding success at time 0 (when the individual enters the study) is assumed to be perfectly correlated with the risk later in life (Wienke 2010). However, this assumption does not necessarily hold, and models accommodating changes in individual latent vital rates may offer an interesting basis to test biological hypotheses. An alternative approach allowing individuals to experience (reversible) changes in latent vital rates could be based on the ontogenetic view of individual differences (Senner et al. 2015). This can be achieved with ‘dynamic frailty’ models (Manda and Meyer 2005, Putter and Van Houwelingen 2014), hidden Markov models (Johnson et al. 2016), Latent Class transition models or mixture models, in which individuals can change latent class over time (e.g. Kaplan 2008). Hidden Markov models are now commonly used in CR studies, but specific applications to change in latent demographic parameters are still rare (Pradel 2005, Cubaynes *et al*. 2010).

### Initial conditions

An overlooked issue in CR studies using multistate models is the issue of ‘initial conditions’. Before estimating the parameters of a model accounting for a stochastic process with dependence between consecutive states (e.g. breeding states), one has to think about how the process was “initialized”. Studies of reproduction necessarily start recording breeding outcomes at the first breeding event (recruitment, or first observed breeding attempt). More generally, studies modeling reproductive outcomes from recruitment onwards (e.g. Cam et al. 1998, Yoccoz et al. 2002) assume that the start of the process generating the observed reproductive states coincides with the start of reproductive life for each individual (Wooldridge 2005). Nevertheless, the process governing the first breeding outcome can be the same as the process generating the subsequent observations in the individual lifetime trajectory (Skrondal and Rabe-hesketh 2014). Such a process can include unobserved determinants of breeding outcome. In dynamic models of reproduction incorporating the effect of past breeding outcome at time *t* on the probability of breeding successfully at time *t+1* (e.g. multistate CR models), the outcome of the first reproductive attempt (at time *t*) is not considered as the realization of a random process, because there is no reproduction at time *t-1*. Nevertheless, failure to incorporate unobserved factors governing breeding success probability at recruitment can translate into overestimation of transition probabilities between subsequent reproductive states (Heckman 1981, Prowse 2012). This is particularly problematic in studies of changes in reproductive costs throughout the lifetime (e.g. Sanz-Aguilar et al. 2008), or of experience-specific variation in breeding outcome (e.g. Nevoux et al. 2007). Interestingly, Sanz-Aguilar et al. (2008) have interpreted evidence of higher reproductive costs of reproduction at recruitment as a consequence of within-cohort mortality selection, with frailer individuals incurring higher reproductive costs than robust ones. The initial conditions problem can be overcome using a joint modeling approach of the processes governing reproductive success at recruitment and subsequent breeding occasions (Skrondal and Rabe-hesketh 2014). If CR data are available from the pre-breeding period, then unobserved and observed determinants of breeding state can be considered simultaneously (e.g. Fay et al. 2016b).

### Inference about individual heterogeneity

Two papers have revived interest in unobserved heterogeneity in demographic parameters in CR studies (Steiner et al. 2010, Orzack et al. 2011). More specifically, these papers have drawn attention to the approaches used to discriminate between hypotheses about sources of variation in CR histories. In CR studies, an influential book by Burnham and Anderson (2002) has promoted the use of multimodel inference such as information criteria to address non-mutually exclusive biological hypotheses about the processes governing mortality, or the arrangement of reproductive states over lifetime trajectories of animals. For example, models accounting for state-dependence in survival or reproduction can be considered (multistate or multievent models; e.g. Sanz-Aguilar et al. 2008), models accounting for unobserved heterogeneity in these demographic parameters too (e.g. Royle 2008, Marzolin et al. 2011), as well as models accounting for both sources of variation in survival and reproduction (e.g. Fay et al. 2016a). This contrasts with approaches based on a single model (namely, state-dependence) and evaluation of the degree of consistency of observed individual CR histories with metrics summarizing key features of histories simulated using parameters estimated with the model in question (Steiner et al. 2010, Orzack et al. 2011).

By definition, variation in individual trajectories simulated using parameters estimated with multistate CR models is not caused by fixed, unobserved heterogeneity between individuals in their demographic parameters (Tuljapurkar et al. 2009, Steiner and Tuljapurkar 2012). The variation in arrangements of states in simulated data stems from the realization of random variables governed by probabilities; the resulting pattern is called ‘dynamic heterogeneity’ (Tuljapurkar et al. 2009), or ‘individual stochasticity’ (Caswell 2009). Several papers have provided evidence that there is a good match between observed and simulated features of individual histories (Steiner et al. 2010, Orzack et al. 2011, Steiner and Tuljapurkar 2012). These studies suggest that stochastic demographic processes have been overlooked in life history studies, and that latent, unobserved heterogeneity in demographic parameters might have been overstated in studies of longitudinal data from animals, whether detection probability is lower than one or not (Cam et al. 2002a, 2013, Steiner et al. 2010, Orzack et al. 2011). From a conceptual viewpoint, these studies attempt to caution biologists against over-interpreting amounts of unobserved individual heterogeneity in demographic parameters (“biologists commonly argue that large differences in fitness components are likely adaptive, resulting from and driving evolution by natural selection” Steiner & Tuljapurkar 2012, Cam et al. 2016). However, they have moved away from one of the dominating statistical inference approaches in the CR area, namely multimodel inference and information criteria (Burnham and Anderson 2002). Current research is addressing the question of whether simulations based on multistate CR models or simply models with state-dependence used for longitudinal data analysis allow discriminating between alternative hypotheses about the processes generating variability in individual histories (Plard et al. 2012, Bonnet and Postma 2016, Cam et al. 2016).

Importantly, proponents of dynamic heterogeneity have overlooked notes of caution from other areas of research also using multistate models for inferences about longitudinal data concerning possible biases in estimates of ‘state-dependence’ (e.g. Heckman 1981, Ahmad 2014). A key issue in discriminating between processes generating variation in individual histories is that a Markov process (i.e. the basis of multistate models) and unobserved individual heterogeneity (for instance a random effect model, Royle 2008) can create similar patterns in arrangements of states along individual trajectories (Ahmad 2014, Authier et al. 2017). This issue has stimulated a large body of work in econometrics (Heckman 1981, Ahmad 2014, Skrondal and Rabe-hesketh 2014, Andriopoulou and Tsakloglou 2015). The hypothesis of a ‘communicating vessels’ phenomenon between sources of variation in CR histories should be considered in wild animal populations, as in econometrics studies (Ahmad 2014, Plum and Ayllón 2015, Cam et al. 2016). Interestingly, several CR studies have hypothesized that their results obtained using multistate models partly reflect heterogeneity between individuals in baseline breeding and survival probability (Cam et al. 1998), or phenotypic within-cohort mortality selection (i.e., the change in the composition of a heterogeneous cohort including individuals with different baseline survival probabilities; Cam et al. 2002a, Barbraud and Weimerskirch 2004, Nevoux et al. 2007, Sanz-Aguilar et al. 2008). That is, they have hypothesized that their result may be caused by unobserved individual heterogeneity, a question now being addressed in studies of senescence (Péron et al. 2010, 2016). This suggests that CR models with both a Markovian structure (for observable, partially observable, or unobservable states) and unobserved individual heterogeneity might perform well with some datasets from wild animal populations (Fay et al. 2016a, b).

## Conclusion

Our review, although not exhaustive, demonstrates that the tremendous advances in CR modeling accomplished over the past 40 years provide investigators with a reliable way to address multiple facets of the process of individual heterogeneity in demographic parameters. Pioneer works by quantitative wildlife biologists focused on individual heterogeneity in recapture or survival probability to avoid biased estimates of population size. The emergence of more general questions such as cause-specific sources of mortality in game- and non-game species (Johnson et al 1986, Koons et al. 2014) and the need for accurate assessment of the impact of global change on the demography of structured populations (Gullett et al. 2014) have triggered collaborations between biologists and statisticians to make efficient use of data, robust inferences about demographic parameters, and achieve an increasing degree of realism in both the sampling and ecological processes handled by CR models. As emphasized by Conroy (2009), the nature of questions that can be addressed nowadays has been greatly expanded to include evolutionary ecology, whose cornerstone is variation in demographic parameters between individuals both within and between populations. The relevance of dealing with individual heterogeneity to study eco-evolutionary processes has placed the topic of individual heterogeneity at the core of many empirical investigations using CR data (Table 1). Provided appropriate sampling design and sufficient data are available, the flexibility of modern CR models now allows assessing reliably the role of individual heterogeneity in ecology and evolutionary processes in the wild.

## Acknowledgements

EC was supported by the Laboratoire d’Excellence LabEx TULIP (ANR-10-LABX-41). OG was supported by ANR-16-CE02-0007. We are grateful to Jim Nichols, Doug Johnson, and Ken Burnham for sharing ideas and material on their early work focusing on the consequences of individual heterogeneity in survival on population dynamics, and the development of CR models accounting for heterogeneity in survival and recapture probability. Evan Cooch provided material on ongoing analyses completing this early work with alternative methods, and Jim Nichols provided helpful comments on an earlier version of this paper. We also thank the three referees for insightful suggestions that improved the paper.

